# Optimisation of a novel method for the production of single-span membrane proteins in *Escherichia coli*

**DOI:** 10.1101/308445

**Authors:** Sarah M. Smith, Kelly L. Walker, Corinne J. Smith, Colin Robinson

**Author notes:** Address for correspondence: Professor Colin Robinson, School of Biosciences, University of Kent, Canterbury, CT2 7NJ.

## Abstract

The large-scale production and isolation of recombinant protein is a central element of the biotechnology industry and many of the products have proved extremely beneficial for therapeutic medicine. *Escherichia coli* is the microorganism of choice for the expression of heterologous proteins for therapeutic application, and a range of high-value proteins have been targeted to the periplasm using the well characterised Sec protein export pathway. More recently, the ability of the second mainstream protein export system, the twin-arginine translocase, to transport fully-folded proteins into the periplasm of not only *E*. coli, but other Gram-negative bacteria, has captured the interest of the biotechnology industry.

In this study, we have used a novel approach to block the export of a heterologous Tat substrate in the later stages of the export process, and thereby generate a single-span membrane protein with the soluble domain positioned on the periplasmic side of the inner membrane. Biochemical and immuno-electron microscopy approaches were used to investigate the export of human growth hormone by the twin-arginine translocase, and the generation of a single span membrane-embedded variant. This is the first time that a *bona-fide* biotechnologically-relevant protein has been exported by this machinery *and* visualised directly in this manner. The data presented here demonstrate a novel method for the production of single-span membrane proteins in *E. coli*.

**Highlights:**

- The Tat translocase has captured the interest of the biotechnology industry
- Biochemical and immuno-EM approaches showed efficient export of hGH by Tat
- A novel approach was used to block export of hGH by Tat in *E. coli*
- We demonstrate a novel method for producing single-span membrane proteins in *E. coli*

## 1.0 Introduction

*Escherichia coli* is responsible for the production of over one third of therapeutic proteins (Walsh, 2010). Conventional strategies for the isolation of these heterologous, recombinant proteins from *E. coli* include, expression of soluble proteins in the cytoplasm, expression as insoluble inclusion bodies or export to the periplasm with subsequent outer membrane rupture to release the periplasmic contents (Pooley *et al.*, 1996). Adoption of the latter approach offers two main advantages: firstly, the oxidizing environment of the periplasm allows disulphide bonds to form, and secondly, rupturing of only the outer membrane of Gram-negative bacteria means there are fewer contaminating cytoplasmic proteins.

The ability of the twin-arginine translocase (hereafter denoted Tat system) to transport fully-folded proteins into the periplasm of Gram-negative bacteria has captured the interest of the biotechnology industry owing to its potential to simplify the downstream processing of high-value, biotherapeutic proteins. Initial studies investigated the ability of the Tat translocase to transport a model heterologous protein on a large scale. These studies showed that Tat could export GFP at levels comparable to conventional methods, and with no large-scale release of cytoplasmic contents (Matos *et al.*, 2012), confirming this machinery’s potential as a viable method for producing heterologous protein.

Most of the Tat system’s substrates are soluble proteins, and most studies on the biotechnological exploitation of the system have similarly focused on soluble high-value protein (Alanen *et al.*, 2015; Browning *et al.*, 2017; Matos *et al.*, 2013). However, the Tat system does have considerable potential for the production of single-span heterologous membrane proteins, particularly those that contain a globular domain located on the periplasmic face of the inner membrane. Two studies have shown that it is possible to ‘stall’ a Tat substrate at the inner membrane by virtue of a substitution mutation in the signal peptidase cleavage site of a TorA signal peptide (Karlsson *et al.*, 2012; Ren *et al.*, 2013). Both studies used biochemical approaches to demonstrate that a Tat substrate remained membrane-anchored with its mature domain exposed at the periplasmic side of the inner membrane. These data reveal a potential for the T at translocase to be used in the production of membrane-bound proteins or for bacterial cell-surface display technologies.

Bacterial surface display involves the presentation of a recombinant protein or peptide on the surface of microorganisms, most common of which is *E. coli*, owing to its ease of genetic manipulation and ability to produce recombinant proteins in high yields. As the name suggests, this presentation technology results in the protein of interest being positioned on the surface of the cell where it is available for interaction with any externally added substrate, which does not have to penetrate the membrane. Implementation of this technique has proven beneficial for numerous applications, including biofuel production and protein library screening (Wu *et al.*, 2008).

A prerequisite of cell surface display in *E. coli* is that the protein of interest is successfully transported across the inner membrane and into the outer membrane; however, incorporation of foreign proteins in the outer membrane can be toxic to the cell; the presence of lipopolysaccharides can sterically hinder interactions between expressed protein and binding-partner, and assembly at cellular appendages (e.g. flagella or pilli) can disrupt assembly of the recombinant protein (Chen & Georgiou, 2002). Therefore, alternative methods are required that avoid the pitfalls of displaying a protein at the outer membrane. Anchored periplasmic expression is one such alternative that involves display of heterologous protein in the periplasm of *E. coli*, which is tethered to the inner membrane by a lipoprotein targeting motif (Mazor *et al.*, 2008).

The vast majority of proteins that are presented using this method are exported out of the cytoplasm in an unfolded state via the Sec translocon (Driessen & Nouwen, 2008). However, this export is reliant on the recombinant protein refolding to a native, biologically active conformation in the periplasm. Use of Tat machinery to traverse the inner membrane avoids this issue since proteins can be transported fully-folded. This, in combination with data showing that a signal peptide mutation in a Tat substrate is able to stall a Tat precursor at the inner membrane (Ren *et al.*, 2013), highlights the potential for Tat machinery in cell surface display technology.

To investigate this further we used biochemical and electron microscopy approaches to analyse the export of a biopharmaceutical, human growth hormone (hGH), which has been shown to be successfully transported by the Tat machinery (Alanen *et al.*, 2015). hGH is a single polypeptide chain of 191 amino acids and is one of the most important hormones in the human body, possessing vital roles in numerous biological processes including cell metabolism and proliferation (Kassem *et al.*, 1993). Recombinant hGH is used therapeutically to treat hGH deficiency and a range of genetic disorders (Sanchez-Ortiga *et al.*, 2012; Spiliotis, 2008; Takeda *et al.*, 2010; Vogt & Emerick, 2015).

Only 2 heterologous proteins have been shown to be anchored in the cell envelope of *E. coli* via the Tat machinery. Maltose binding protein (MBP) was anchored at the inner membrane via the addition of a 22-residue C-terminal transmembrane helix (from the native Tat substrate, HybO) (Karlsson *et al.*, 2012). The same study also detected scFv13 (human antibody fragment specific for ß-galactosidase) at the inner membrane when a FLAG tag was positioned between the signal peptide and the scFv domain.

In this study we have sought to achieve stable expression of a non-cleavable Tat precursor at the inner membrane of *E. coli* without the addition of amino acid residues to the protein, and we have used immunogold electron microscopy to precisely localise the target proteins for the first time.

Direct immunogold labelling revealed that overexpressed, membrane-targeted hGH exhibits a uniform distribution in the inner membrane of *E. coli*. The data confirm that this type approach can be used for the display of globular protein domains on the periplasmic face of the inner membrane.

## 2.0 Materials & Methods

### 2.1 Export assays of hGH in *E. coli* cells

*E. coli* cells (transformed with a plasmid containing either WT or mutant precursor hGH (Alanen *et al.*, 2015)), were cultured in 1L LB media and induced with 1 mM arabinose (or 1 mM IPTG) and cultured for 2 hours. At given intervals (0 hr, 1 hr, 1 hr 15 min, 1 hr 30 min, 1 hr 45 min and 2 hrs), 10 ml of culture was harvested and the cell pellet was normalized against an optical density of 600 nm (e.g. for an optical density of 0.6 at 600 nm, the cell pellet was resuspended in 0.6 ml of disruption buffer [100 mM Tris-acetate pH 8.2, 500 mM sucrose and 5 mM EDTA]).

### 2.2 Fractionation of *E. coli* cells and detection of hGH protein

*E. coli* cells were fractionated into periplasmic, cytoplasmic and membrane fractions using the lysozyme/cold osmotic shock method (Randall & Hardy, 1986). The resulting fractions were analysed by sodium dodecyl sulphate polyacrylamide gel electrophoresis (SDS PAGE).

Once the periplasmic, cytosolic and membrane samples had been analysed using SDS-PAGE the proteins were transferred from acrylamide gels to PVDF membranes via semi-dry Western Blotting apparatus. The membranes were blocked overnight in a solution of 5% (w/v) dried milk in PBS-T and then incubated with primary anti-hGH antibody (1:20000, TBS/Tween [0.24% Tris (w/v), 2.5% NaCl (w/v) and 2% Tween-20 (v/v), pH 8.4]) for 1 hour before washing. Membranes were incubated with anti-rabbit-HRP conjugate (Promega, WI, USA) for 1 hr at RT, and subsequently washed. Finally, immunoreactive bands were detected using ECL^™^ detection reagents according to the manufacturer’s instructions. X-ray films were developed using an AGFA Curix 60 automatic developer as directed by the manufacturer’s instructions.

### 2.3 Preparation of *E. coli* cells for visualisation by electron microscopy

Chemical fixation of *E. coli* cells: *E. coli* cells (overexpressing either WT or mutant precursor hGH) were resuspended in aldehyde fixative (0.25% glutaraldehyde/4% formaldehyde) and low melting temperature agarose at 1:1 volume ratio. Agarose-enrobed cells were then harvested and incubated on ice until the agarose had set. Cells were diced into 1 mm^3^ pieces using a clean razor blade and resuspended in fresh aldehyde fixative for overnight fixation at 4 °C. Chemically fixed cells were washed for 1-2 hrs in 0.15 M sodium cacodylate/HCl buffer pH 7.4 at 4 °C to remove aldehyde fixative. The reaction was then quenched for 1 hr by washing in cacodylate buffer containing 0.1M glycine. Contrast was imparted to the cells by incubation in 1% tannic acid (TA) for 1 hr at 4°C, followed by washing in H_2_O for a second staining of 1% uranyl acetate (aq) for 1 hr at 4°C. *E. coli* cells were dehydrated in an ethanol series (70-100%) for a period of 2hrs for subsequent resin embedding (London Resin Company Ltd.; Berkshire, UK).

### 2.4 Ultrathin sectioning of resin blocks for imaging by transmission electron microscopy

Ultrathin (50 nm) sections were cut (using an ultramicrotome [Ultracut E, Reichert-Jung; Vienna, Austria]) and transferred to 200-mesh carbon-coated copper grids (Agar Scientific, Stansted Essex) for immunolabelling of hGH (section 2.5) and subsequent transmission-EM imaging.

### 2.5 Immunolabelling of *E. coli* cells to detect hGH in the inner membrane

Using the touching-drop method (Rubinstein, 2007), ultrathin sections were blocked via incubation in TBS/Tween buffer (0.24% Tris (w/v), 2.5% NaCl (w/v) and 2% Tween-20 (v/v)) pH 8.4 containing 1% BSA (w/v) and 4% normal goat sera (v/v) (Abcam Plc, Cambridge UK) for 30 mins at room temperature (RT). hGH protein was labelled by incubation with primary anti-hGH antibody (rabbit polyclonal) at 1:20000 dilution (TBS/Tween +1% BSA buffer) for 2hrs at RT. Sections were then washed with TBS/Tween +1% BSA at RT, and then incubated with 10nm gold-conjugated secondary antibody (Goat anti-rabbit IgG pre-adsorbed, Abcam Plc) for 1 hr at RT. Finally, sections were washed in TBS/Tween (+1% BSA) and dH_2_O and allowed to air dry before insertion into the EM.

### 2.6 Transmission electron microscopy of ultrathin *E. coli sections*

Ultrathin sections of *E. coli* cells were imaged under a 200 kV JEOL 2010F transmission electron microscope with a field emission gun electron source, operating a Gatan Ultrascan^™^ 4000 CCD camera with a pixel size of 14 μm.

### 2.7 Quantification of immunogold labelling

A random sampling approach was used to analyse the distribution of gold within the immunolabelled *E. coli* sections, as in (Smith *et al.*, 2017). An identical Chi-squared analysis procedure (as in (Smith *et al.*, 2017)) was performed on samples of *E. coli* cells overexpressing either WT or mutant precursor hGH, with the null hypothesis (of no difference in immunolabelling between hGH-overexpressing cells and wild-type *E. coli* cells not overexpressing any recombinant protein) being rejected (p<0.005).

## 3.0 Results

### 3.1 TorA-hGH is exported efficiently by the Tat system

hGH was expressed with a N-terminal TorA signal peptide that contains several features typical of a Tat signal peptide (hereafter referred to as TorA-hGH). These include an N-terminal domain, a hydrophobic core region and a polar C-terminal domain ending with an Ala-Xaa-Ala consensus motif (Figure 1). It is this signal peptide that targets hGH, and other proteins, to the inner membrane of *E. coli* for efficient export into the periplasm by the Tat machinery (Ren *et al.*, 2013).

**Figure 1.**
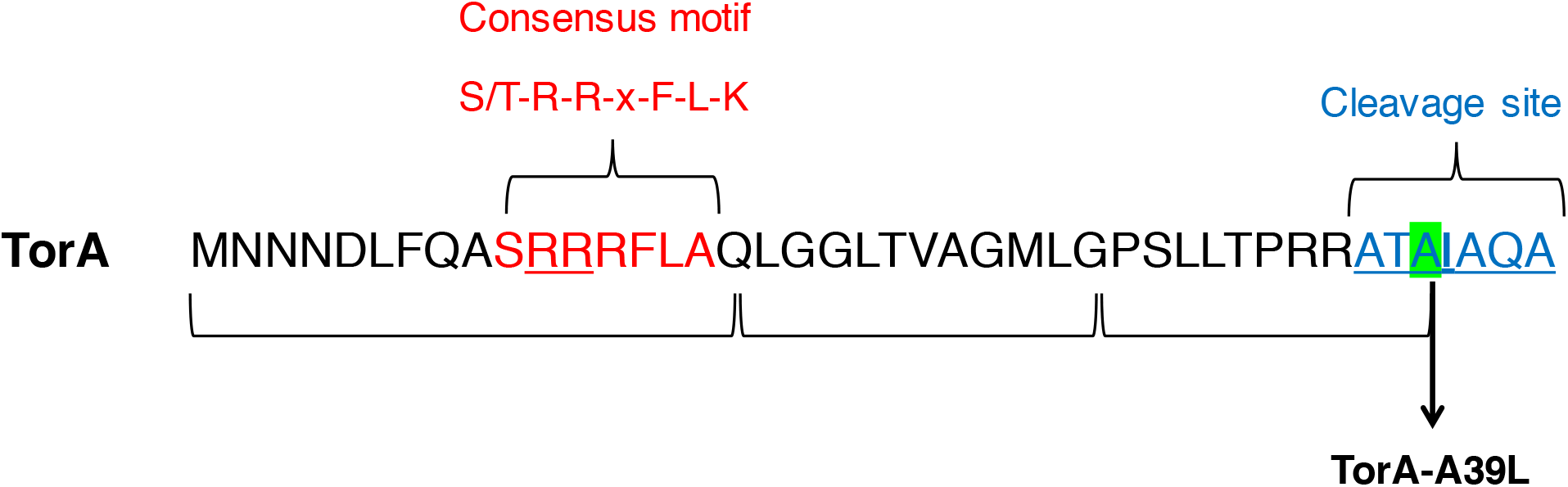
Structure of the TorA signal peptide and the TorA-A39L variant. Primary structure of the 42-residue signal peptide of *E. coli* TorA. The consensus motif is highlighted in red (twin-arginine motif underlined), and the peptidase cleavage site is highlighted in blue. The terminal, alanine residue at position 39 (highlighted in green), was substituted by leucine to generate the TorA-A39L variant.

(Alanen *et al.*, 2015) showed that hGH can be exported by Tat and subsequently acquire its disulphide bonds in the periplasm. The aim of this study was to gain a more detailed insight into the export of this biopharmaceutical in *E. coli*, and in particular to directly visualise the location of the exported protein. We initially used standard export assays to examine the export of TorA-hGH in wild-type MC4100 *E. coli* cells expressing native levels of Tat machinery. At certain time points, up until 2 hours after the induction of the plasmid, cells were fractionated to generate cytoplasm (C), periplasm (P) and membrane & insoluble (MI) samples which were analysed by SDS-PAGE and immunoblotting.

Immunoblotting against hGH protein showed two forms of hGH are produced in whole cell fractions: a 27 kDa precursor protein and 22 kDa mature protein (Figures 2, 3 and 4). Some non-specific bands are also observed.

**Figure 2.**
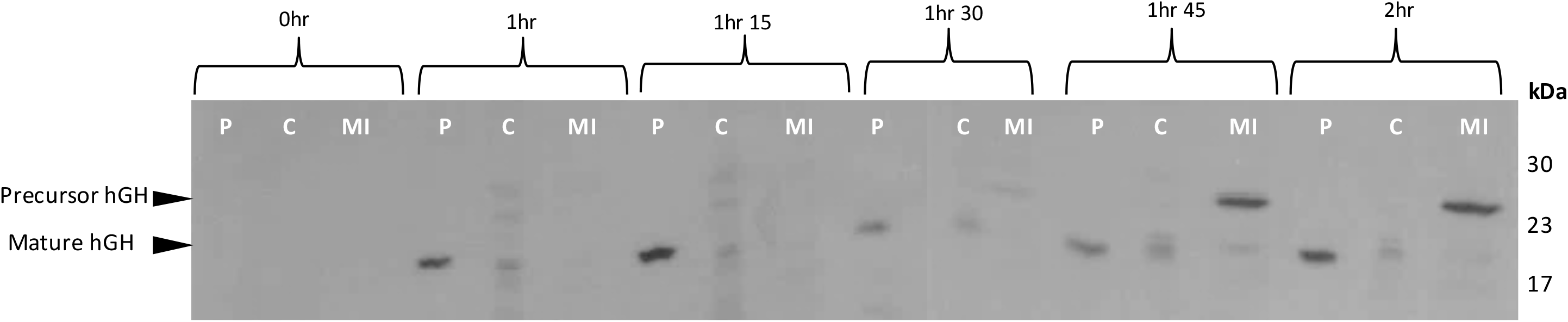
Export assay of Tat substrate (TorA-hGH) into the periplasm of *E. coli.* Western Blot to detect the presence of hGH in *E. coli*. Post-induction with 1 mM IPTG *E.coli* cells overexpressing hGH fused to a TorA signal peptide (TorA-hGH) were grown at 37°C for 2 hours. During this time, cells were periodically harvested, normalised for OD_600_ = 10 and subsequently fractionated to periplasm (P), cytoplasm (C) and membrane/insoluble fractions (MI) fractions. Each fraction was examined for the presence of hGH by immunoblotting with anti-hGH antibody. Results show there is a good level of export from as early as 1 hour after induction, shown by the presence of mature hGH (22 kDa) in the periplasm. At each time point there is a lack of pre-cursor (27kDa) TorA-hGH (i.e. Tat substrate) in the cytoplasmic fractions, indicating the substrate is either degraded or forming inclusion bodies in the cell. From 1 hr 45min onwards there is a consistent level of precursor TorA-hGH in the membrane fraction of the *E. coli* cells suggesting to the formation of inclusion bodies.

**Figure 3.**
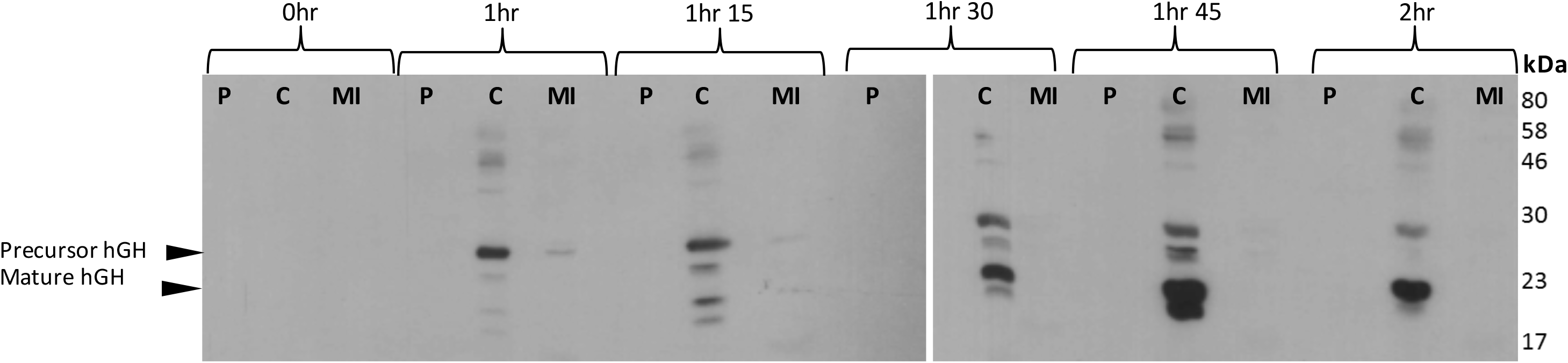
Export assay of TorA-hGH into the periplasm of *E. coli* cells lacking expression of Tat machinery. Western Blot to detect the presence of hGH in *E. coli*. Post-induction with 1 mM IPTG *E.coli* cells overexpressing hGH fused to a TorA signal peptide (TorA-hGH) and lacking expression of Tat machinery i.e. Δ*tat* were grown at 37°C for 2 hours. During this time, cells were periodically harvested, normalised for OD_600_ = 10 and subsequently fractionated to periplasm (P), cytoplasm (C) and membrane/insoluble fractions (MI) fractions. Each fraction was examined for the presence of hGH by immunoblotting with anti-hGH antibody. Results show that mature hGH (22 kDa) is present in the cytoplasm from very early on (1 hr). Precursor hGH (27 kDa) expression increases at 1 hr 30, and there is no hGH present in the MI fraction at all time points tested.

**Figure 4.**
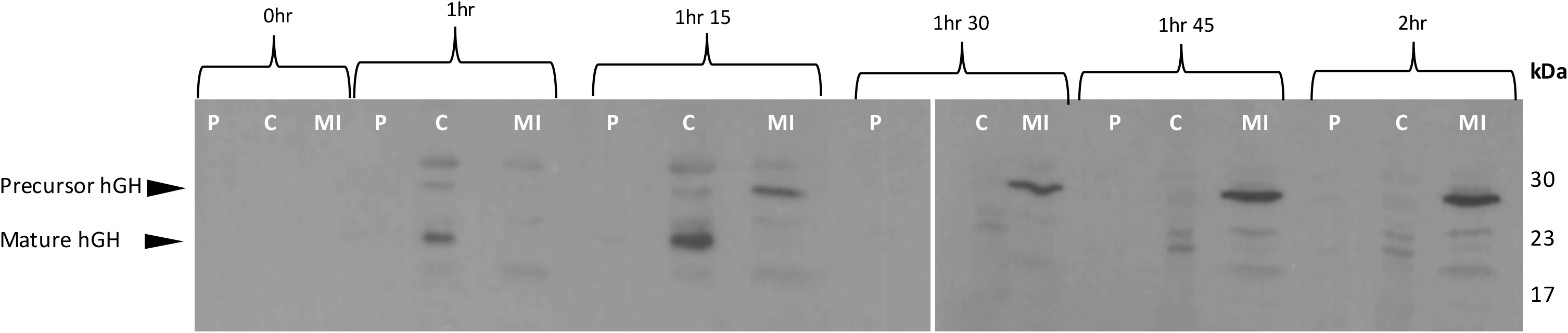
Export assay of a non-cleavable Tat substrate (TorA(A39L)-hGH) into the periplasm of *E. coli.* Western Blot to detect the presence of non-cleavable hGH in *E. coli*. Post-induction with 1 mM IPTG *E.coli* cells overexpressing hGH fused to a non-cleavable TorA signal peptide (TorA-hGH) were grown at 37°C for 2 hours. During this time, cells were periodically harvested, normalised for OD_600_ = 10 and subsequently fractionated to periplasm (P), cytoplasm (C) and membrane/insoluble (MI) fractions. Each fraction was examined for the presence of hGH by immunoblotting with anti-hGH antibody. A lack of hGH protein in the periplasm of any time point confirms that the Tat substrate is non-cleavable and thus remains in the inner membrane fraction. hGH is degraded in the cytoplasm from as early as 1 hr indicated by the mature sized hGH (22 kDa). This is degraded completely by 1 hr 30 mins. From 1 hr 15 min onwards there is a consistent level of precursor TorA-hGH (27 kDa) in the membrane fraction of the *E. coli*.

Export assays demonstrated that Tat-mediated transport of TorA-hGH occurs soon after induction since, after 1 hour, mature hGH is present in the periplasm. There is a minimal amount of precursor protein present in the cytoplasmic fraction, which has also been observed in previous analyses of Tat-dependent export of TorA-hGH (Ren *et al.*, 2013). Therefore, the lack of cytoplasmic precursor protein in this study could be due to the quick turnover of the substrate from the cytoplasm to the periplasm, or due to proteolytic cleavage of the precursor protein (Figure 2). Quick turnover of the precursor hGH would also explain the lack and/or minimal amount of precursor hGH present at the membrane up until 1hr 30. Only at later time points (1hr 45 and 2hr) do we see a substantial increase in precursor at the membrane, which is consistent with increased production of hGH precursor/Tat substrate.

To confirm that the export of TorA-hGH is via Tat, an identical export assay was performed in *E. coli* cells lacking Tat machinery (Δ*tatABCDE*; hereafter denoted Δ*tat*). The lack of mature hGH in the periplasm of these control cells, at any time point, confirms Tat-dependent export of hGH into the periplasm in wild type *E. coli* (Figure 3).

### 3.2 Substitution of the −1 position of the signal peptide blocks maturation of TorA-hGH

It has previously been shown that the nature of the amino acid side chain at the −1 position of the Tat signal peptide (i.e., the last residue of the signal peptide) is essential for efficient maturation of a Tat precursor protein (Ren *et al.*, 2013). The maturation enzyme, leader peptidase, will only tolerate a short chain residue such as alanine at this position. Substitution of the alanine residue at the −1 position of the signal peptide by leucine completely blocked maturation of a native Tat substrate, YedY, with the protein remaining membrane-bound and its mature domain exposed to the periplasm (Ren *et al.*, 2013).

In order to test whether the same mutation blocks maturation of hGH - a biotechnologically relevant, heterologous Tat substrate - export assays were conducted as in section 3.1, with the −1 alanine residue in the TorA signal peptide of TorA-hGH substituted with leucine (generating TorA-A39L-hGH).

Consistent with the findings of (Alanen *et al.*, 2015; Ren *et al.*, 2013), the precursor protein was not processed to any significant degree and is found almost exclusively in the membrane fraction as precursor protein (Figure 4, 1hr 15 time point onwards). As seen with the wild-type precursor, mature hGH was detected in the cytoplasm from an early time point (1hr), which most likely represents proteolytic cleavage of the signal peptide, as observed previously (Alanen *et al.*, 2015)

### 3.3 Direct visualisation of a biotherapeutic, hGH, in *E. coli*

#### 3.3.1 Visualisation of TorA-hGH

Harnessing of industrial microorganisms, such as *E. coli*, for the production of desired products is reliant on the efficient performance of these ‘cell factories’. As such, numerous factors serve as important indicators of platform fitness including: protein localisation and distribution and cellular morphology. Previous studies that have analysed the export of biotechnologically-relevant proteins by the Tat machinery have primarily used biochemical approaches (Karlsson *et al.*, 2012; Ren *et al.*, 2013), yet direct visualisation of the microorganism, in this case *E. coli*, could provide novel insight into the export of such substrates. In this study, we analysed the location of TorA-hGH to test whether the exported protein is indeed randomly distributed in the periplasm, and focused particular attention on the non-cleavable A39L-hGH to determine whether the protein is uniformly distributed in the inner membrane.

To do this, *E. coli* cells overexpressing TorA-hGH or TorA(A39L)-hGH were fixed and stained. 2D sections of individual *E. coli* cells were then cut, immunolabelled and finally examined for the presence of gold particles. Use of antibodies specific to hGH avoids the use of (potentially large) affinity tags, and thus permits direct protein detection. Gold particles were classified according to their location, and for the present study, a gold particle found within 25 nm of the inner membrane was defined as being located at the inner membrane. Our aim was to achieve unambiguous identification of the hGH protein in the inner membrane through immunogold labelling of *E. coli* cells overexpressing TorA-hGH or TorA(A39L)-hGH. To do this, the level of non-specific antibody labelling was assessed through comparison of hGH-overexpressing cells with MC4100 *E. coli* cells that did not overexpress any recombinant protein.

Figure 5 shows representative images of TorA-hGH-overexpressing cells that were immunogold-labelled with anti-hGH. These cells exhibit a pattern of gold labelling that is predominantly around the periphery of the cell. This was confirmed at increased magnification, as in Figures 5 (bottom row), where the gold particles could be seen to reside within 25 nm of the inner membrane of the cell wall. Given that the bulk of the protein has been shown to be present in periplasmic fractions using export assays (Figure 2), we conclude that these particles are indeed located in the periplasm of the cells. A smaller number of particles were present in the cytoplasm. Magnified images of these cells, with close-ups of individual gold particles can be also seen in Supplementary Figure 1.

**Figure 5.**
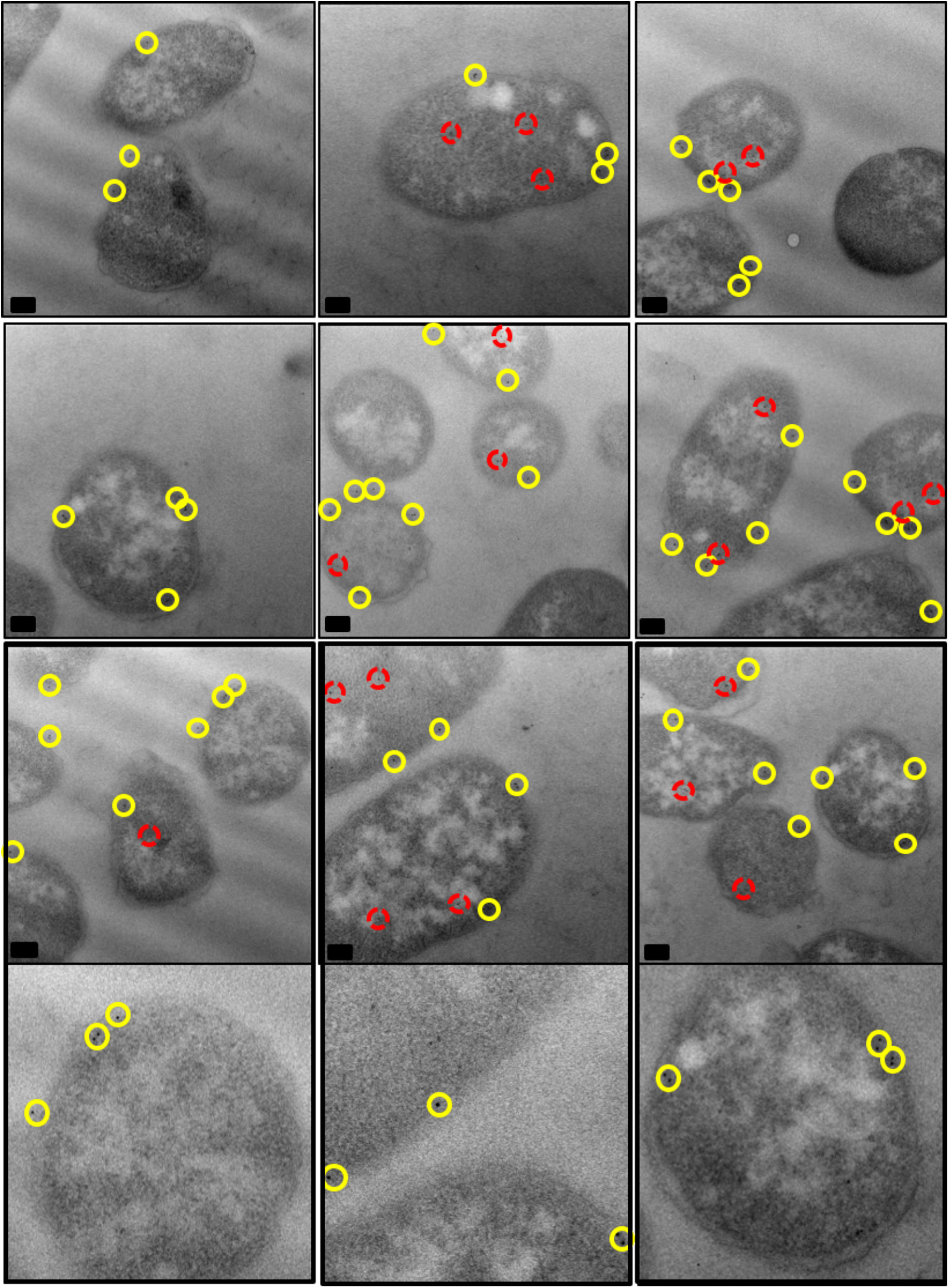
Electron micrographs of *E. coli* cells, overexpressing TorA-hGH (WT precursor), immunogold-labelled following primary antibody detection against hGH protein. Ultrathin sections of *E. coli* cells overexpressing TorA-hGH (WT precursor) were immunolabelled using a polyclonal antibody raised against hGH (shown in rows 1-3, with row 4 showing close-ups of individual gold particles from row 3). At 1 hr 45 min after induction, hGH was found to exhibit a random distribution in the inner membrane (yellow circles) and was also present in the cytoplasm (red circles). Images were taken on a JEOL 2010F at 15,000X magnification. Scale bar = 200 nm.

Analysis of *E. coli* cells that did not express TorA-hGH (representative images shown in Supplementary Figure 2), showed the presence of only a few gold particles in the cytoplasmic and membranous regions. These data indicate that the antibody is able to detect hGH with a high degree of specificity. As further controls, TorA-hGH-expressing cells were immunolabelled with the primary antibody omitted (Supplementary Figure 3, A and B), and TEM analysis showed that these cells lacked any gold binding confirming that non-specific binding seen in *E. coli* cells not overexpressing any recombinant protein was attributable to the primary antibody. Finally, the same cell type was immunolabelled with a gold-conjugated secondary antibody directed towards a different animal species (Supplementary Figure 3, C and D). TEM analysis showed only unlabeled *E. coli* cells, confirming that the cells do not have a non-specific attraction for gold particles.

To quantitate the degree of specific labeling, raw gold counts were collected from 200 randomly sampled, immunolabelled *E. coli* with or without the overexpression of TorA-hGH, and the gold particles were assigned to either the cytoplasmic or inner membrane/ periplasm compartments. Quantification of the gold particles (Figure 6) shows that the average number of gold particles per compartment in TorA-hGH-overexpressing cells was 1.92 per cell (±0.41 gold) in the cytoplasm and 4.29 per cell (±0.37 gold) in the inner membrane, whereas *E. coli* cells lacking overexpression of TorA-hGH bound 0.86 per cell in the cytoplasm (±0.19 gold) and 0.46 (±0.15 gold) in the membrane, respectively. The labelling of membrane-bound TorA-hGH is thus highly specific, with the inner membrane/ periplasm of TorA-hGH-overexpressing cells containing 9.42-fold more gold particles than the same membrane in *E. coli* cells that lack overexpression of TorA-hGH. To confirm a statistical independence in immunogold labelling between these cells at the cytoplasm and inner membrane, *χ*^2^ analysis of the raw gold count data was conducted. For a total *χ*^2^ value of 111.2 and 1 degree of freedom P was <0.005. This confirms that the gold labelling distributions between the two cell types are significantly different.

**Figure 6.**
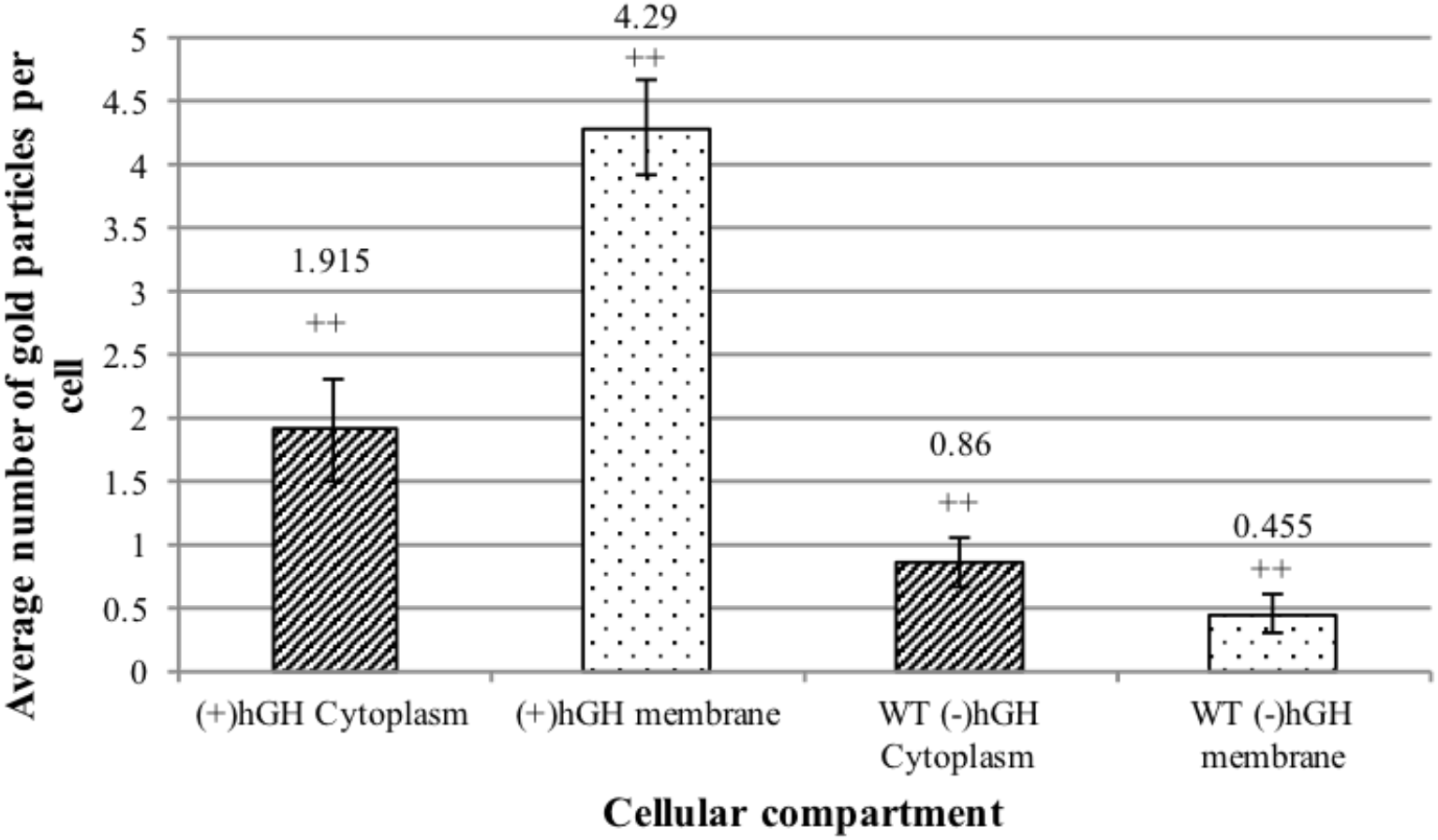
Quantitative analysis of raw gold counts of *E. coli* cells immunogold labelled with primary antibody detection against TorA-hGH protein. *E. coli* cells overexpressing TorA-hGH were immunolabelled using a primary antibody against the hGH protein. Controls were *E. coli* cells that lacked expression of hGH protein (labelled –hGH). Raw gold counts were taken from 200 randomly imaged *E. coli* from 2 separate resin blocks. Gold was assigned to either ‘membrane + periplasm’ (labelled membrane) or cytoplasm compartment. The approximate numbers of gold particles at the cytoplasm and membrane fraction of each cell were calculated for each cell type. There was 2.2X more gold in the cytoplasm (line-patterned) of hGH expressing cells versus those lacking hGH expression. There was 9.43× more gold in the membrane (dotted pattern) of hGH expressing cells versus those lacking hGH expression. Error bars: CI of 2× SE. The labelling of hGH between the two cell types is statistically significant (indicated by ++).

Analysis of the distribution of gold particles in the cytoplasm shows the presence of 1.92 and 0.86 particles in the TorA-hGH-overexpressing and non-hGH expressing *E. coli* cells, respectively. Since the number of gold particles in *E. coli* cells that lack overexpression of hGH represents non-specific binding, and assuming that the level of non-specific binding in the two cell types is the same, this indicates that TorA-hGH-overexpressing cells contain an average of 1 gold particle per cell in the cytoplasm, which is considerably lower than the 3.8 gold particles per cell in the inner membrane/ periplasm region. This difference is consistent with the amount of hGH in the cytoplasmic fraction compared to the membranous compartment of *E. coli* determined by export assays, which revealed that at 1hr 45 mins after induction, there is considerably more hGH present in the membrane fraction compared to the cytoplasm (Figure 2). Our immunolabelling approach thus provides highly specific detection of hGH in both the cytoplasmic and membrane fractions of *E. coli*.

Immunogold labelling of TorA-hGH showed this protein is randomly distributed within the *E. coli* periplasm and cytoplasm, with no evidence for a preferential localisation at the poles or elsewhere (Figure 5, yellow and red circles, respectively). This is the first time that a heterologous Tat substrate has been directly visualised after export to the periplasm, and these observations are consistent with the efficient production of hGH in our *E. coli* cells.

#### 3.3.2 Visualisation of non-cleavable TorA-hGH

To investigate whether a similar distribution and/or localisation occurs for the mutated TorA-hGH construct, (TorA(A39L)-hGH), we immunogold labelled *E. coli* cells overexpressing TorA(A39L)-hGH to detect the presence of hGH. Analysis of the localisation and distribution of gold particles showed that, similar to the wild-type precursor (TorA-hGH, section 3.3.1), TorA(A39L)-hGH exhibits a random distribution in the cytoplasm (red circles) and a uniform distribution in the inner membrane (yellow circles) of the *E. coli* cell (Figure 7). This was confirmed at higher magnifications, as in Figures 7 (bottom row) and Supplementary Figure 4. Therefore, in addition to biochemical export assays (Figure 4) which showed a lack of TorA(A39L)-hGH in the periplasm, these EM data provide direct evidence that an unprocessed Tat precursor remains membrane-associated despite being unable to undergo cleavage by signal peptidases in the periplasm.

**Figure 7.**
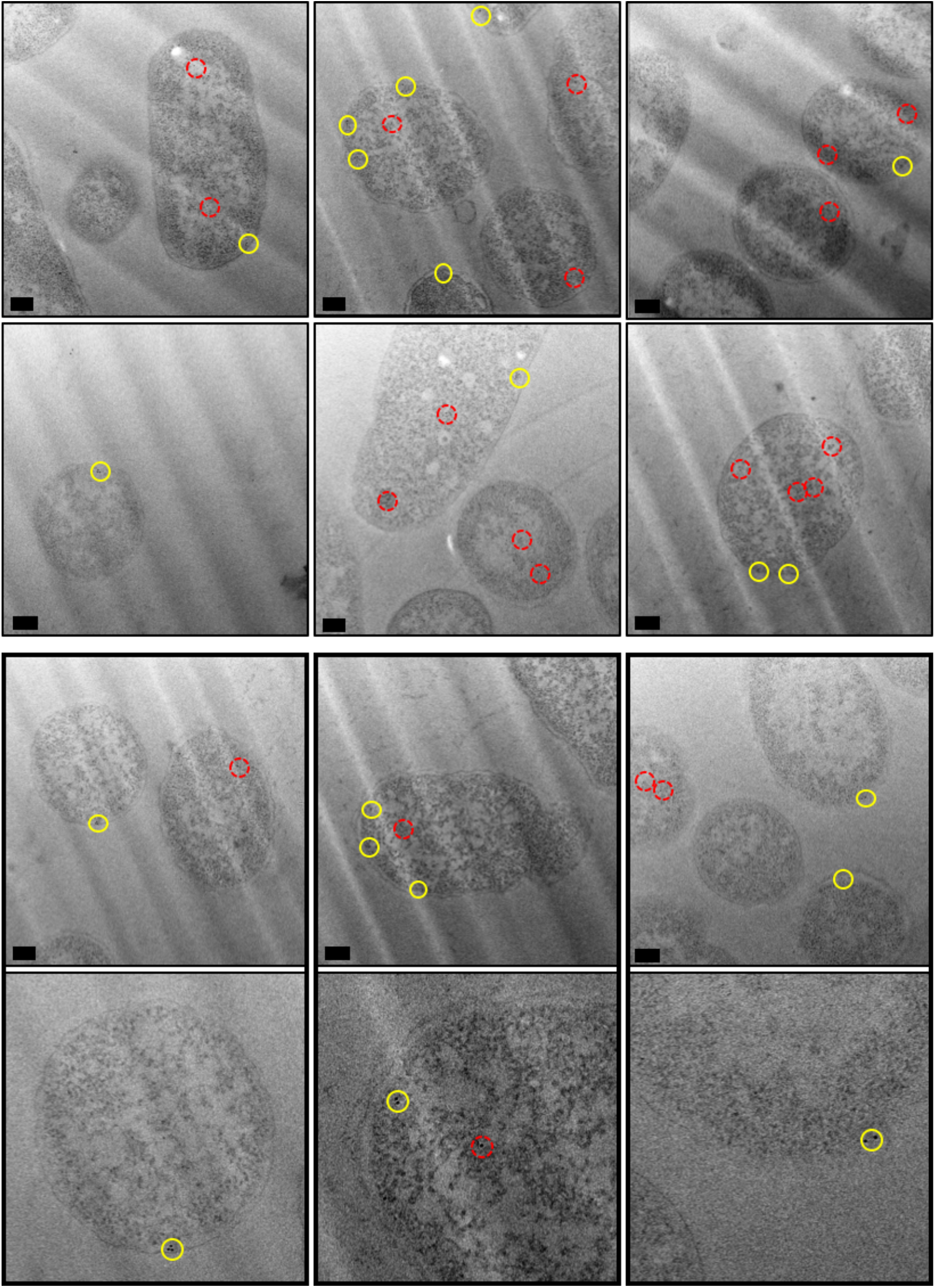
Electron micrographs of *E. coli* cells, overexpressing TorA-hGH (mutant precursor), immunogold-labelled following primary antibody detection against hGH protein. Ultrathin sections of *E. coli* cells overexpressing TorA-hGH (mutant precursor) were immunolabelled using a polyclonal antibody raised against hGH (shown in rows 1-3, with row 4 showing close-ups of individual gold particles from row 3). At 1 hr 45 min after induction, hGH was found to exhibit a random distribution in the inner membrane (yellow circles) and was also present in the cytoplasm (red circles). Images were taken on a JEOL 2010F at 15,000X magnification. Scale bar = 200 nm.

## 4.0 Discussion

The twin-arginine translocase offers potential for the biotechnology industry owing to its ability to transport fully-folded proteins into the periplasm of *E. coli*. Initial studies confirmed the capability of the Tat system to transport a model heterologous protein to a high yield (Matos *et al.*, 2012), and more recent data has shown that substrates are not limited to GFP with Tat being able to transport hGH, scFv and interferon α2b (Alanen *et al.*, 2015; Browning *et al.*, 2017). Furthermore, it has also been shown that it is possible to anchor a Tat substrate in the inner membrane of *E. coli* with the mature domain of the protein facing the periplasmic side of the membrane (Karlsson *et al.*, 2012; Ren *et al.*, 2013). Combined, these data reveal a novel potential for the Tat system in producing single-span membrane proteins in *E. coli*.

Anchoring of heterologous Tat substrates in the inner membrane has previously occurred via the addition of extra amino acid residues to the protein sequence (Karlsson *et al.*, 2012). However, this study has shown that only a single substitution mutation in the signal peptidase cleavage site of a TorA signal peptide, fused to hGH, resulted in the precursor protein being “stalled” at the inner membrane, with no alterations to the mature protein. The previous studies by (Karlsson *et al.*, 2012) and (Ren *et al.*, 2013) showed that cleavage by leader peptidase occurs only after transfer of the mature protein to the periplasm, and given that hGH is exported by Tat in an extremely efficient manner, it is to be expected that the mature hGH protein will be similarly positioned on the periplasmic face of the inner membrane.

In summary, a combination of biochemical export assays and imaging approaches were used in this study to investigate the temporal expression of TorA-hGH and TorA(A39L)-hGH in *E. coli* and to determine whether they were transported via Tat machinery. Our data demonstrate the Tat-dependent transport of TorA-hGH into the periplasm and most importantly, the stable expression of TorA(A39L)-hGH at the inner membrane, revealing a potential novel methodology for the production of single-span membrane proteins in *E. coli* via the Tat machinery.

## Acknowledgements

Support for K.L.W as part of the Bioprocessing Research Industry Club (BRIC), a partnership between the BBSRC, EPSRC and a consortium of biotechnology companies, is gratefully acknowledged. We thank Ian Hands-Portman (Imaging suite and Advanced Bioimaging Research Technology Platform, University of Warwick) for support and access to the Jeol 2010F and 2011 TEMs (Welcome Trust grant 055663/Z/98/Z).

## References

Alanen, H. I., Walker, K. L., Lourdes Velez Suberbie, M., Matos, C. F. R. O., Bönisch, S., Freedman, R. B., Keshavarz-Moore, E., Ruddock, L. W. & Robinson, C. (2015) Efficient export of human growth hormone, interferon α2b and antibody fragments to the periplasm by the Escherichia coli Tat pathway in the absence of prior disulfide bond formation. Biochimica et Biophysica Acta (BBA) - Molecular Cell Research, 1853 (3): 756–763.

Browning, D. F., Richards, K. L., Peswani, A. R., Roobol, J., Busby Stephen, J. W. & Robinson, C. (2017) Escherichia coli “TatExpress” strains super-secrete human growth hormone into the bacterial periplasm by the Tat pathway. Biotechnology and Bioengineering, 114 (12): 2828–2836.

Chen, W. & Georgiou, G. (2002) Cell-Surface Display of Heterologous Proteins: From High-Throughput Screening to Environmental Applications Biotechnology and Bioengineering, 79 (5): 496–503.

Driessen, A. J. M. & Nouwen, N. (2008) Protein Translocation Across the Bacterial Cytoplasmic Membrane. Annual Review of Biochemistry, 77 (1): 643–667.

Karlsson, A. J., Lim, H.-K., Xu, H., Rocco, M. A., Bratkowski, M. A., Ke, A. & DeLisa, M. P. (2012) Engineering Antibody Fitness and Function Using Membrane-Anchored Display of Correctly Folded Proteins. Journal of Molecular Biology, 416 (1): 94–107.

Kassem, M., Blum, W., Ristelli, J., Mosekilde, L. & Eriksen, E. (1993) Growth hormone stimulates proliferation and differentiation of normal human osteoblast-like cells in vitro. Calcified Tissue International, 52 (3): 222–226.

Matos, C., Branston, S., Albinak, A., Dhanoya, A, Freedman, R, Keshavarz-Moore, E. & Robinson, C. (2012) High-yield export of a native heterologous protein to the periplasm by the tat translocation pathway in Escherichia coli. Biotechnology & Bioengineering, 109 (10): 2533–2542.

Matos, C. F. R. O., Robinson, C., Alanen Heli, I., Prus, P., Uchida, Y., Ruddock, L. W., Freedman, R. B. & Keshavarz-Moore, E. (2013) Efficient export of prefolded, disulfide-bonded recombinant proteins to the periplasm by the Tat pathway in Escherichia coli CyDisCo strains. Biotechnology Progress, 30 (2): 281–290.

Mazor, Y., Van Blarcom, T., Iverson, B. L. & Georgiou, G. (2008) E-clonal antibodies: selection of full-length IgG antibodies using bacterial periplasmic display. Nature Protocols, 3 1766.

Pooley, H. M., Merchante, R. & Karamata, D. (1996) Overall Protein Content and Induced Enzyme Components of the Periplasm of Bacillus subtilis. Microbial Drug Resistance, 2 (1): 9–15.

Randall, L. L. & Hardy, S. J. S. (1986) Correlation of competence for export with lack of tertiary structure of the mature species: A study in vivo of maltose-binding protein in E. coli. Cell, 46 (6): 921–928.

Ren, C., Patel, R. & Robinson, C. (2013) Exclusively membrane-inserted state of an uncleavable Tat precursor suggests lateral transfer into the bilayer from the translocon. FEBS Journal, 280 (14): 3354–3364.

Rubinstein, J. L. (2007) Structural analysis of membrane protein complexes by single particle electron microscopy. Methods, 41 (4): 409–416.

Sanchez-Ortiga, R., Klibanski, A. & Tritos, N. (2012) Effects of recombinant human growth hormone therapy in adults with Prader-Willi syndrome: a meta-analysis. Clinical Endocrinology, 77 (1): 86–93.

Smith, S. M., Yarwood, A., Fleck, R. A., Robinson, C. & Smith, C. J. (2017) TatA complexes exhibit a marked change in organisation in response to expression of the TatBC complex. Biochemical Journal, 474 (9): 1495–1508.

Spiliotis, B. (2008) Recombinant human growth hormone in the treatment of Turner syndrome. Therapeutics and Clinical Risk Management, 4 (6): 1177–1183.

Takeda, A., Cooper, K., Bird, A., Baxter, L., Frampton, G., Gospodarevskaya, E., Welch, K. & Bryant, J. (2010) Recombinant human growth hormone for the treatment of growth disorders in children: a systematic review and economic evaluation. Health Technology Assessment, 14 (42): 1–209.

Vogt, K. & Emerick, J. (2015) Growth hormone therapy in adults with Prader-Willi syndrome. Diseases, 3 56–67.

Walsh, G. (2010) Post-translational modifications of protein biopharmaceuticals. Drug Discovery Today, 15 (17-18): 773–780.

Wu, C. H., Mulchandani, A. & Chen, W. (2008) Versatile microbial surface-display for environmental remediation and biofuels production. Trends in Microbiology, 16 (4): 181–188.

